# Temporal dynamics of *Candida albicans* morphogenesis and gene expression reveals distinctions between in vitro and in vivo filamentation

**DOI:** 10.1101/2024.02.20.581211

**Authors:** Rohan S. Wakade, Melanie Wellington, Damian J. Krysan

## Abstract

*Candida albicans* is a common human fungal pathogen that is also a commensal of the oral cavity and gastrointestinal tract. *C. albicans* pathogenesis is linked to its transition from budding yeast to filamentous morphologies including hyphae and pseudohyphae. The centrality of this virulence trait to *C. albicans* pathobiology has resulted in extensive characterization a wide range factors associated with filamentation with a strong focus on transcriptional regulation. The vast majority of these experiments have used in vitro conditions to induce the yeast-to-filament transition. Taking advantage of in vivo approaches to quantitatively characterize both morphology and gene expression during filamentation during mammalian infection, we have investigated the dynamics of these two aspects of filamentation in vivo and compared them to in vitro filament induction with “host-like” tissue culture media supplemented with serum at mammalian body temperature. Although filamentation shares many common features in the two conditions, we have found two significant differences. First, alternative carbon metabolism genes are expressed early during in vitro filamentation and late in vivo, suggesting significant differences in glucose availability. Second, *C. albicans* begins a hyphae-to-yeast transition after 4hr incubation while we find little evidence of hyphae-to-yeast transition in vivo up to 24hr post infection. We show that the low rate of in vivo hyphae-to-yeast transition is likely due to very low expression of *PES1*, a key driver of lateral yeast in vitro, and that heterologous expression of *PES1* is sufficient to trigger lateral yeast formation in vivo.

**Importance:** *Candida albicans* filamentation is correlated with virulence and is an intensively studied aspect of *C. albicans biology*. The vast majority of studies on *C. albicans* filamentation are based on in vitro induction of hyphae and pseudohyphae. Here we used an in vivo filamentation assay and in vivo expression profiling to compare the tempo of morphogenesis and gene expression between in vitro and in vivo filamentation. Although the hyphal gene expression profile is induced rapidly in both conditions, it remains stably expressed over a 12hr time course in vivo while it peaks after 4hr in vitro and is reduced. This reduced hyphal gene expression in vitro correlates with reduced hyphae and increased hyphae-to-yeast transition. In contrast, there is little evidence of evidence of hyphae-to-yeast transition in vivo.

## Introduction

*Candida albicans* is a component of the human mycobiome that also causes disease in both immunocompetent and immunocompromised patients (1, 2). The transition of *C. albicans* from harmless commensal to invasive pathogen is associated with a morphological switch from budding yeast to filaments comprised of both hyphae and pseudohyphae (3). In general, *C. albicans* strains and mutants that show low rates of filamentation are more fit in the commensal setting and less fit during invasive infection (4, 5). Over the years, filamentation has been one of the most intensively studied *C. albicans* virulence traits (3). The vast majority of these studies used one of a number of in vitro conditions to generate filaments. Recently, we developed an in vivo imaging approach for the analysis of *C. albicans* filamentation during infection. In this approach, *C. albicans* is inoculated directly into the subdermal ear tissue of the mouse (6). Anatomically, this compartment is stromal tissue beneath an epithelium, in this case skin epithelium, and shares many features with stromal tissue beneath colonized epithelium of the mouth and GI tract. We have used this approach to identify the transcription factor network that regulates filamentation in vivo (7) as well as to characterize the filamentation of clinical isolates and protein kinase mutants (8, 9). These studies have revealed a number of important differences in the genes required for filamentation in vivo compared to in vitro. For example, the cAMP-Protein Kinase A pathway is absolutely required for in vitro filamentation in all inducing- media but is dispensable for filamentation in vivo (9).

The first goal of this work was to characterize the temporal dynamics of filamentous morphogenesis and associated gene expression in vivo beginning at early time points using the imaging assay. We coupled this with in vivo expression analysis using Nanostring nCounter technology to profile the expression of a set of 186 environmentally responsive genes over the same time periods (7, 9). This set of genes includes 57 (30%) hyphae-associated genes (7). Our previous Nanostring-based in vivo expression profiling of *C. albicans* infection of ear tissue, kidney tissue and oral tissue at single time points has shown both similarities and differences within these niches (9). Furthermore, expression profiles of in vitro hyphal induction also show similarities and differences to in vivo expression.

To further explore the similarities and differences between filamentation under in vivo and in vitro conditions, we followed morphogenesis and gene expression over an identical 12hr time course of in vitro and in vivo filamentation. Nanostring nCounter technology was used to analyze gene expression for two primary reasons. First, genome-wide RNA-seq methods for the direct analysis of *C. albicans* gene expression in infected tissue have not been developed. Although a gene-enrichment strategy has been reported (10), its application to time course analysis was cost prohibitive. Second, we were most interested in the temporal dynamics of a set of well-studied hyphae-associated environmentally responsive genes; therefore, genome-wide characterization of expression was not necessary for our purposes (11). As such, this approach limits the conclusions we can make about global patterns of gene expression.

A single in vitro filament-inducing condition was used and was chosen because it is generally used in the field to approximate a host-like environment. Specifically, the tissue culture medium RPMI 1640 supplemented with 10% bovine calf serum (BCS) was used and incubations were performed at mammalian body temperature (37°C). Previous microarray-based expression profiling of the time course for in vitro *C. albicans* filamentation used rich medium (YPD or YEPD) supplemented with 10% BCS (12). It has become well-established that the specific filament-inducing medium has a significant effect on both the regulatory pathways and gene expression patterns involved in *C. albicans* filamentation (13). Indeed, those considerations motivated our study of gene expression pattern in vivo.

To date, we have performed hundreds of in vivo imaging assays with WT cells, the majority of which examined filamentation at 24hr. In vitro, hypha begin to form lateral yeast cells after ∼4hr of induction; however, we have rarely observed lateral yeast cell formation in vivo. Therefore, the second goal of this study was to explore the mechanistic basis underlying this distinction between in vitro and in vivo morphogenesis. As detailed below, our data suggest that *C. albicans* begin a hyphae-to-yeast transition that is correlated with a reduction in the expression of hyphae-associated genes after ∼4hr induction (14) and that this is likely to involve *PES1*, a gene known to drive lateral yeast formation in vitro (15). In vivo, *C. albicans* cells maintain expression of hyphae-associated dreams and have very low expression of *PES1* throughout the time course, a result that explains why we find little evidence of the hyphae-to-yeast transition in vivo. Consistent with this model, we show that expression of *PES1* from a strong, heterologous promoter is sufficient to drive lateral yeast formation in vivo.

## Results

### In vivo filamentation is initiated rapidly and reaches steady state twelve hours after infection

Stationary phase *C. albicans* SN250 labeled with NEON were injected into the subdermal tissue of mice ears and imaged at 1hr, 2hr, 4hr, 8hr and 12hr post infection (Fig. 1A). Germ tube-like cells with short filaments are initially observed in 60% of cells 1hr post-infection (Fig. 1B&C). The percentage of cells that have a filamentous morphology increase slightly until 4hr at which a steady state ratio of filamentous cells to yeast cells was reached. Previously reported data at 24hr indicate that there is little change in this ratio between 12hr and 24hr. The length of the filaments increases over the first few hours of the time course (Fig. 1D) and reaches a steady state at 8hr. The median length remains constant between 8hr and 12hr and comparison to previously reported data for 24hr indicates that the median length of the population remains relatively stable to that time point.

**Figure 1.**
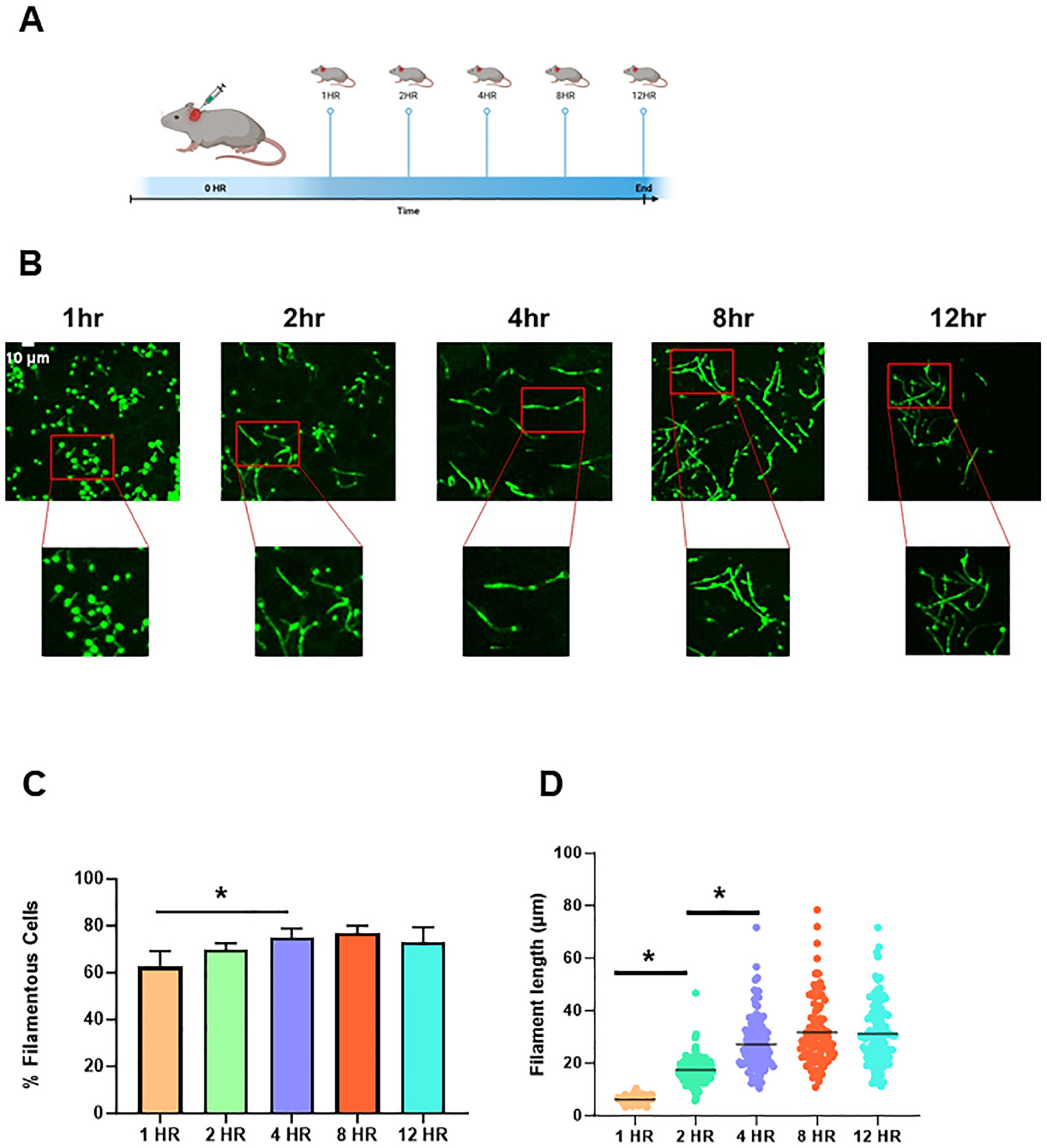
Time course of in vivo *C. albicans* filamentation. **A**. Diagram of time course experiment. **B.** Representative 2-D fields from confocal microscopy of NEON-labeled SN250 strain in ear tissue at the indicated time points after infection. **C**. Quantification of percent filamentous cells at the indicated time points. Asterisks indicated significant difference between time points (p < 0.05 *, ANOVA with Tukey’s method of multiple comparison correction). **D.** The length of filaments at the indicated time points. Asterisks indicate significant difference between the groups (p < 0.05 *, Mann Whitney test).

The same time course experiment was conducted for in vitro filamentation by inoculating stationary phase cells into RPMI 1640 with 10% BCS at 37°C and fixing cells at each time point. As expected, germ tubes are also initially observed in vitro after 1hr (Fig. 2A&B) but the proportion of cells that have formed a germ tube is significantly less in vitro relative to in vivo conditions after 1hr (60% in vivo and 20% in vitro). The percentage of filamentous cells peaks at 4hr. In contrast to in vivo conditions, the percentage of filamentous cells then declines by ∼25% between 4hr and 12hr (p <0.05). Similar to in vivo filamentation, the median length of the filaments increases over the first 4h to a steady state (Fig. 2C). These data indicate that in vitro filamentation in RPMIS is slightly delayed relative to in vivo conditions and that the ratio of filaments to yeast begins to decline after a peak at 4hr.

**Figure 2.**
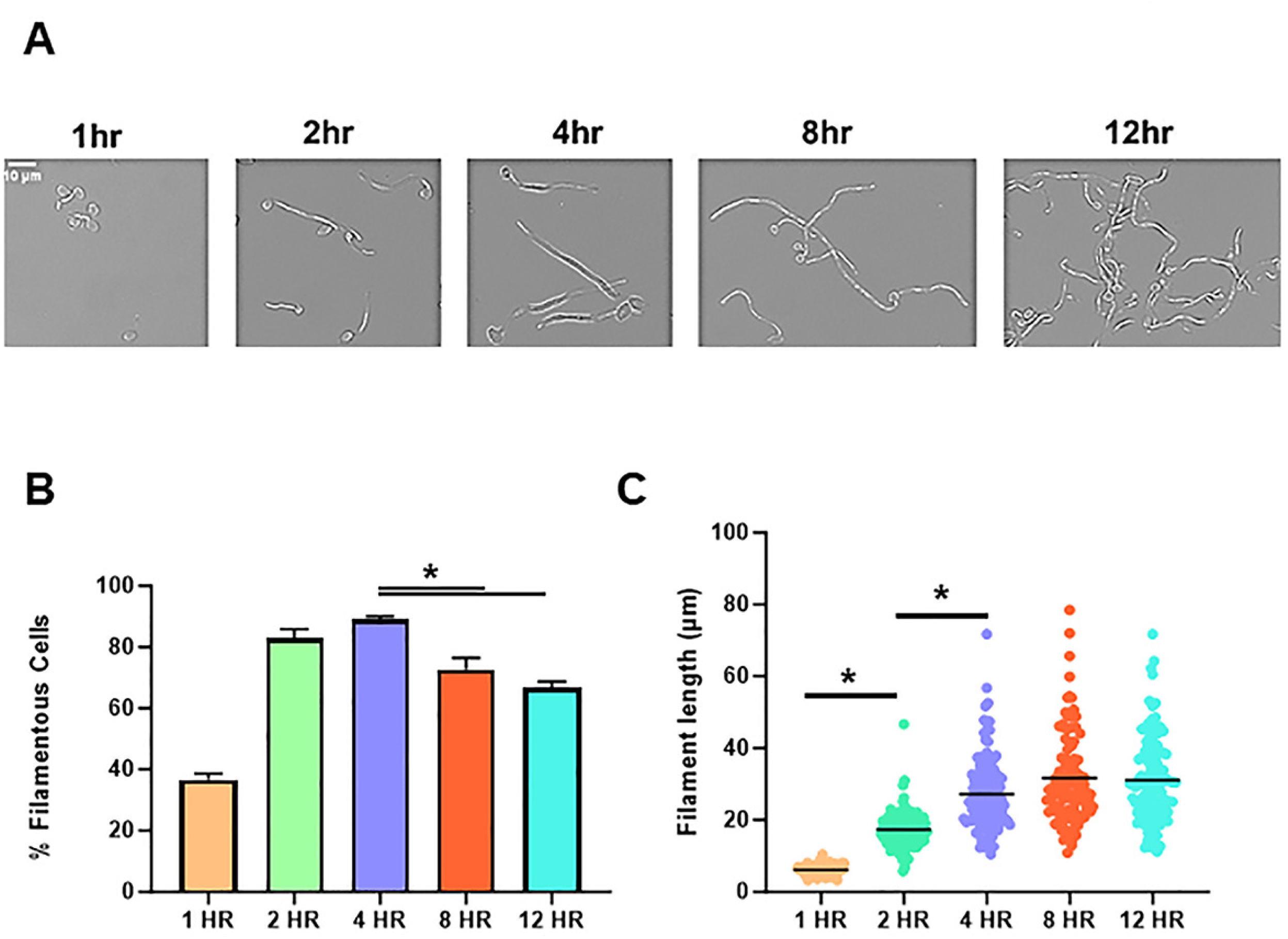
Time course of in vivo *C. albicans* filamentation. **A**. Representative bright field images of cell morphology following hyphal induction in RPMI tissue culture medium supplemented with 10% bovine calf serum at 37°C for the indicated times. **B**. Quantification of percent filamentous cells at the indicated time points. Asterisk indicated significant difference between time points (p < 0.05 *, ANOVA with Tukey’s method of multiple comparison correction). **C.** The length of filaments at the indicated time points. Asterisks indicate significant difference between the groups (p < 0.05 *, Mann Whitney test).

### Expression profiles of in vivo and in vitro filaments evolve over the time course

Nanostring nCounter methods were used to characterize the expression of 186 genes and compared them to the yeast inoculum at 1, 2, 4, 8, and 12hr post-infection (Table S1) as well as post filament induction in vitro (Table S2). The raw, normalized, and processed data are provided in Tables S1&S2 for each time point. Differentially expressed genes were defined as those with ± 2-fold change in expression with an FDR <0.1 as determined by the Benjamini- Yeuketil procedure when compared to expression in the yeast phase inoculum which was grown overnight at 30°C in rich medium (yeast peptone with 2% dextrose, YPD). The time course of transcriptional changes is summarized by volcano plots in Fig. 3A and 3B for in vivo and in vitro conditions, respectively. The total number of differentially expressed genes at each time point is indicated in Fig. 3A and 3B. We generated Venn diagrams to compare the genes upregulated under in vitro conditions to in vivo at each time point as a way to assess the similarity and differences between the two conditions at the same time point (Fig. 4A-E). At 1hr, the set of genes induced by a statistically significant amount is low because of relatively high variability (Table S1); this variability resolves by 2hr. As expected, the set of upregulated genes common to both in vivo and in vitro conditions at 1hr includes regulators of hyphae morphogenesis (*BRG1*, *CPH1/2*, *TEC1*, & *UME6*) as well as hyphae-associated cell wall genes (*ALS1* & *HWP1*); see Table S1 graphs below (Fig. 5&6).

**Figure 3.**
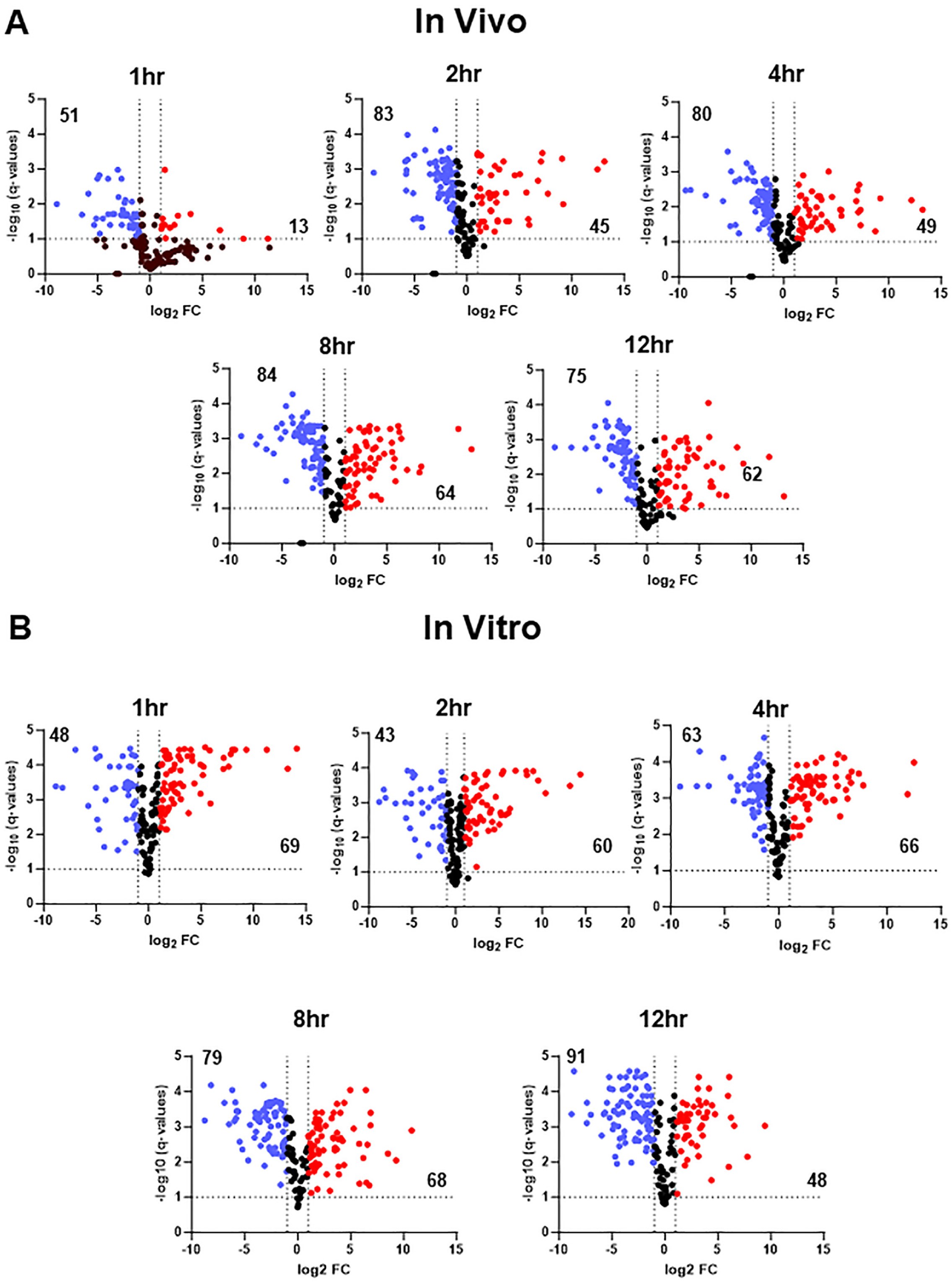
Nanostring analysis of gene expression over time for in vivo and in vitro filamentation. Volcano plots showing genes with significant (log_2_ ± 1; FDR<0.1, Benjamini method) increase (red dots) or decrease (blue dots) at the indicated time points for in vivo filamentation (**A**) and in vitro filamentation (**B**). Expression is normalized to yeast phase cells used to infect mice or inoculate in vitro cultures. The numbers in the two quadrants indicate total number of differentially expressed genes for that region of the plot.

**Figure 4.**
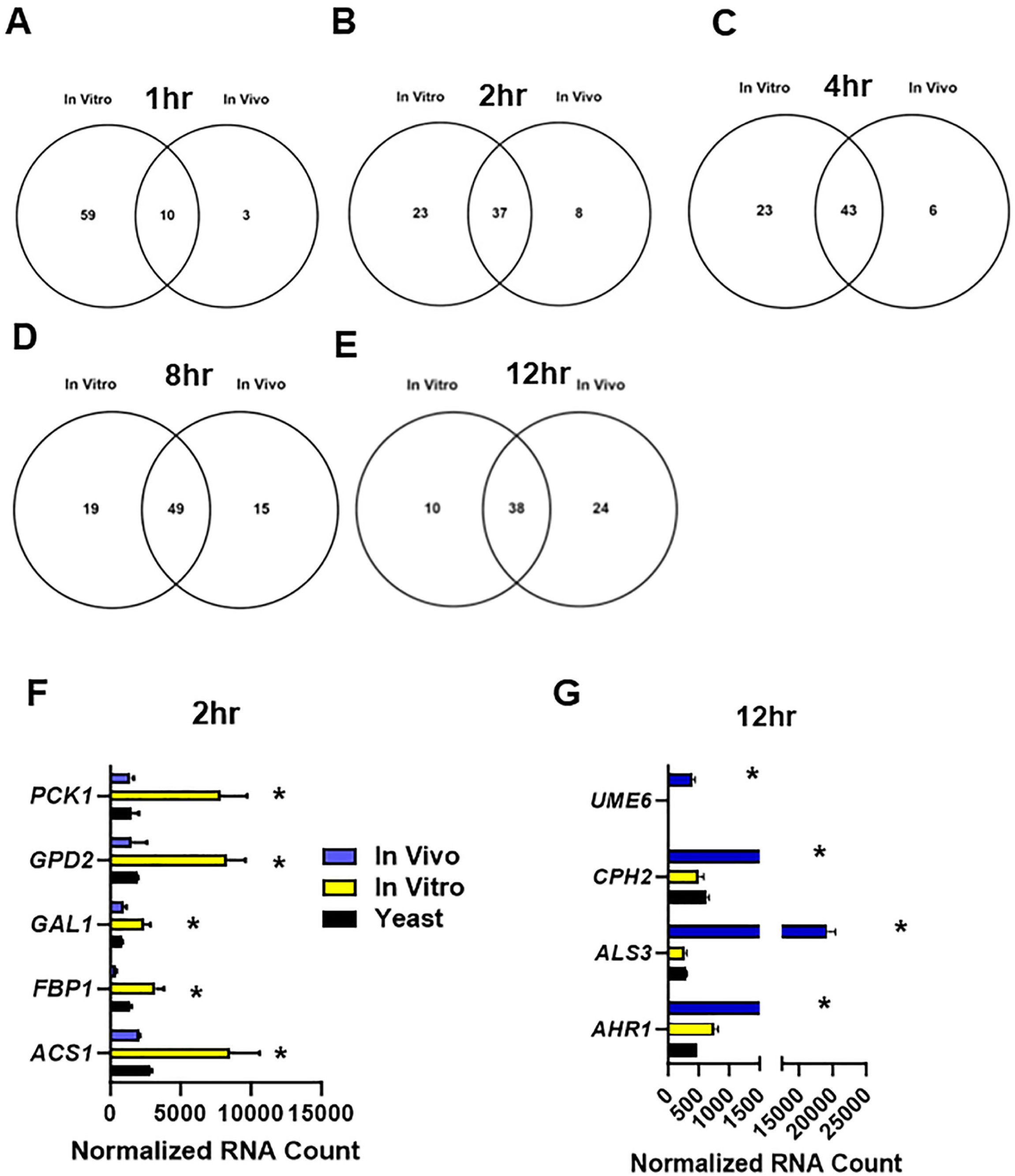
Distinct patterns of upregulated genes during in vitro and in vivo filamentation. Venn diagrams comparing the profiles of upregulated genes in vitro and in vivo at 1hr (**A**); 2hr (**B**); 4hr (**C**); 8hr (**D**) and 12hr (**E**). **F**. Set of alternative carbon metabolism genes upregulated at the 2hr time point during in vitro induction relative to yeast inoculum (* indicates FDR <0.1) but not significantly different from inoculum in vivo. **G.** Set of hypha-associated genes upregulated in vivo at the 12hr time point relative to yeast inoculum (* indicates FDR <0.1) but not significantly different from inoculum in vitro. Bars indicate normalized mRNA counts from three independent experiments with error bars indicating standard deviation.

**Figure 5.**
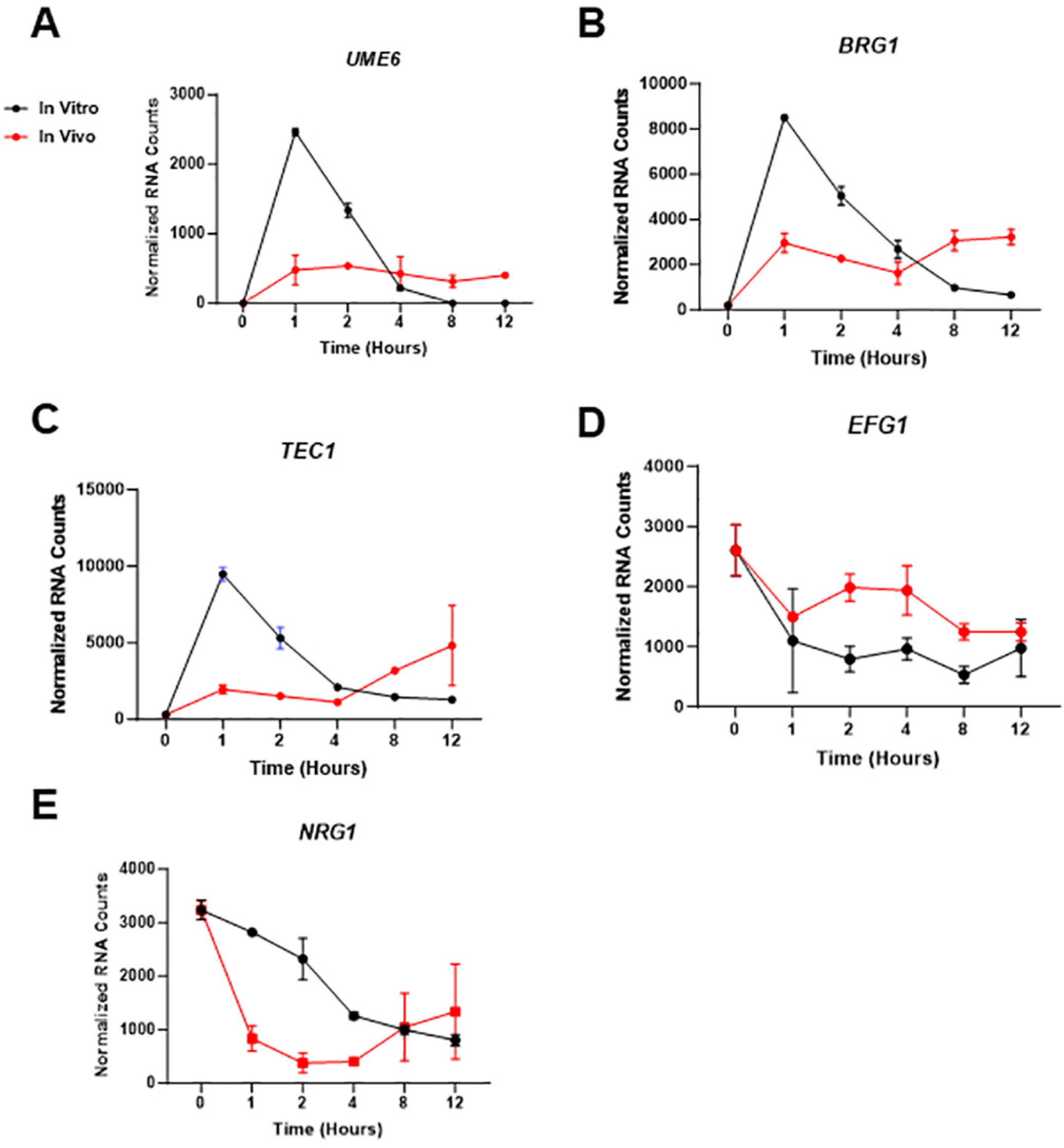
Comparison the dynamics of expression for hyphae-associated transcriptional regulators during in vitro and in vivo filamentation. **A**. Normalized mRNA counts for the expression of *UME6* (**A**), *BRG1* (**B**), *TEC1* (**C**), *EFG1* (**D**), and *NRG1* (**E**) at the indicated time points. Bars indicate normalized mRNA counts from three independent experiments with error bars indicating standard deviation.

At early time points (Fig. 4A-C), the majority of genes upregulated in vivo are also upregulated in vitro. However, a large proportion of genes are uniquely upregulated in vitro. For example, one-third of the genes upregulated in vitro at 2hr were not upregulated in vivo. The set uniquely upregulated in vitro is enriched for carbon metabolism genes (FDR 0.01, Benjamini- Yeuktiel: *PCK1*, *GPD2*, *GAL1*, *ACS1*, *FBP1*, Fig. 4F); *PCK1* and *FBP1* are involved in gluconeogenesis and *ACS1* is induced by low glucose (16), suggesting that glucose availability may be distinct between in vitro and in vivo conditions at early time points. Further supporting that possibility, *AOX2*, which is induced by low glucose conditions (17), was strongly upregulated at the 1hr (59-fold increase relative to yeast, FDR = 0.005) and 2hr (22- fold- increase relative to yeast, FDR = 0.006) time points in vitro, while *AOX2* expression is below the level of detection in vivo (Table S1). At later time points in vivo, *AOX2* (12 hr, 69-fold increase relative to yeast, FDR = 0.02) and *PCK1* were upregulated (12hr, 4-fold increase relative to yeast, FDR = 0.05), suggesting that the cells begin to depend more on non-glucose carbon sources as the time course progresses.

At the 12hr time point, more genes are uniquely upregulated in vivo compared to in vitro (Fig. 4E). The set of 24 genes upregulated only in vivo at the 12hr time point is enriched for zinc (FDR: 2e^-4^; *ZAP1*, *ZRT1* and *PRA1*) and iron (FDR: 2e^-3^; *HAP3*, *FTR1*, *RBT5*, and *PGA7*) homeostasis (18, 19). Interestingly, filamentous growth associated genes are also enriched in this set (FDR: 3e^-4^) including transcriptional regulators of filamentous growth *UME6*, *AHR1*, and *CPH2* and the hyphae-associated cell wall gene *ALS3* (Fig. 4G). We were particularly struck by the fact that *UME6* expression was no longer upregulated after 12hr of induction in vitro. Indeed, *UME6* expression had returned to the near background levels of expression by 8hr of in vitro induction and was not different than in yeast phase (Fig. 5A). *UME6* expression is required for the maintenance of hyphal elongation both in vitro and in vivo (7, 20). The decline in the expression of *UME6* after 8hr of in vitro hyphal induction suggests that the hyphal transcriptional program is reduced late in the time course whereas in vivo the activity of this program appears to be maintained longer.

### The expression of hyphae-induced genes decreases over time in vitro and correlates with reduced filaments at those time points

To further evaluate the possibility that the hyphal morphogenic and transcriptional program was waning after 4hr in vitro but not in vivo, we compared the expression of transcription factors that positively regulate filamentation in vitro and in vivo. In addition to *UME6*, the hyphae-induced transcription factors (TFs) *BRG1* and *TEC1* are upregulated after 1hr exposure to RPMIS and 1hr post-infection in vivo (Fig. 5A-C). In vitro, the expression of all three TFs peaks and then falls. In contrast, in vivo expression of *UME6*, *BRG1*, and *TEC1* shows a slower slope of initial induction and then maintains relatively stable levels throughout the time course. The hyphae-associated TFs *CPH1* and *CPH2* also follow the same pattern of expression but are not as strongly induced (Tables S1 and S2). *EFG1* is a critical regulator of the hyphal transcriptional profile under many conditions (21). Interestingly, and in contrast to the other three TFs, *EFG1* expression is reduced relative to yeast phase both in vitro and in vivo (Fig. 5D). Finally, expression of the repressor of filamentation *NRG1* (22, 23) is also downregulated under both conditions, although this downregulation occurs slightly more rapidly in vivo (Fig. 5E).

Consistent with the temporal pattern for the expression of hyphae-induced TFs in vitro, the expression of hyphae-associated target genes *ALS3*, *ECE1*, *HWP1*, *HYR1* and *IHD1* show the same peak and decline over time. Once again, the tempo for the expression of these genes in vivo is distinct with a more gradual increase followed by relatively stable expression over the time course (Fig. 6A-E). We also examined the expression of the yeast-specific cell wall protein *YWP1* (24) over the time course to see if the reduced expression of hyphae-associated genes was accompanied by a corresponding increase the expression of a yeast associated gene. As shown in Fig. 6F, the expression of *YWP1* declines in both in vitro and in vivo conditions and remains low relative to yeast with a slight increase in expression under both conditions after a nadir at 4hr. Thus, despite the reduction in the expression of hyphae-associated TFs and other hyphae associated gene at later time points in vitro, expression of the *YWP1* remains low at those same time points relative to yeast phase cells. Overall, the decreased expression of hyphae-associated genes at later time points correlates with the decrease in the proportion of filaments after a peak between at approximately 4hr. The expression of the same genes in vivo is stable over the same time period and also correlates with the stable extent of filamentation over the same time points in vivo.

**Figure 6.**
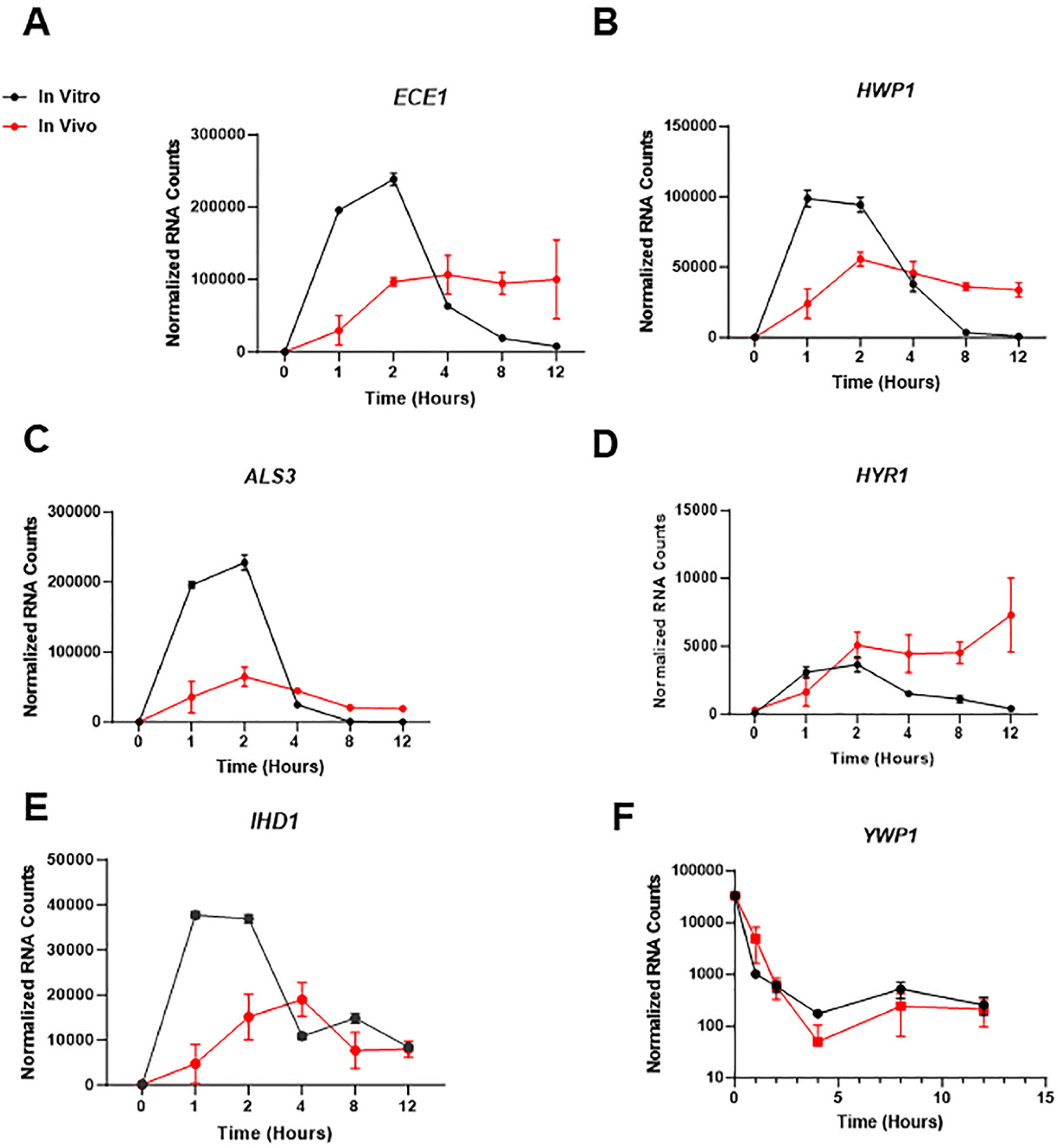
Comparison the dynamics of expression for hyphae- and yeast associated genes during in vitro and in vivo filamentation. **A**. Normalized mRNA counts for the expression of *ECE1* (**A**), *HWP1* (**B**), *ALS3* (**C**), *HYR1* (**D**), *IHD1* (**E**) and *YWP1* (**F**) at the indicated time points. Bars indicate normalized mRNA counts from three independent experiments with error bars indicating standard deviation.

### Over a 24hr time course in vitro filaments generate lateral yeast but in vivo filaments do not

In vitro, yeast cells begin to emerge at subapical cell bodies in a process termed, the hyphae-to-yeast transition. In our previous large-scale screen comparing in vivo filamentation between TF mutants and WT in one-to-one competition experiments (over 150 replicates of WT, ref. 7), we observed very few lateral yeasts on in vivo filaments at the 24hr point. Although the hyphae-to-yeast transition remains a relatively understudied aspect of *C. albicans* morphogenesis (14), lateral yeast formation is linked to the expression of *PES1* (15). Pes1 is a pescadillo protein that is required for yeast growth and lateral yeast formation as well as biofilm dispersion (25). We, therefore, compared the expression of *PES1* during in vitro and in vivo filamentation. In vivo, *PES1* expression drops dramatically to below or at background for the majority of the time course (Fig. 7A). In contrast, *PES1* expression initially increases during in vitro induction and then falls to essentially background by 12hr post-induction. As reported by Shen et al., over-expression of *PES1* from the *TET*- promoter leads to increased lateral yeast cell formation during a variety of in vitro hyphae induction conditions (15). We, therefore, tested if this was the case for RPMIS using the same strain generated by Shen et al (15); because previous work indicated that lateral yeast formation tends to increase with time, we extended this experiment to 24hr (14). Consistent with previous reports, the *tetO-PES1* strain generated more lateral yeast than a congenic strain with *PES1* expressed (pWT-*PES1*) from its native promoter at all time points (Fig. 7B).

**Figure 7.**
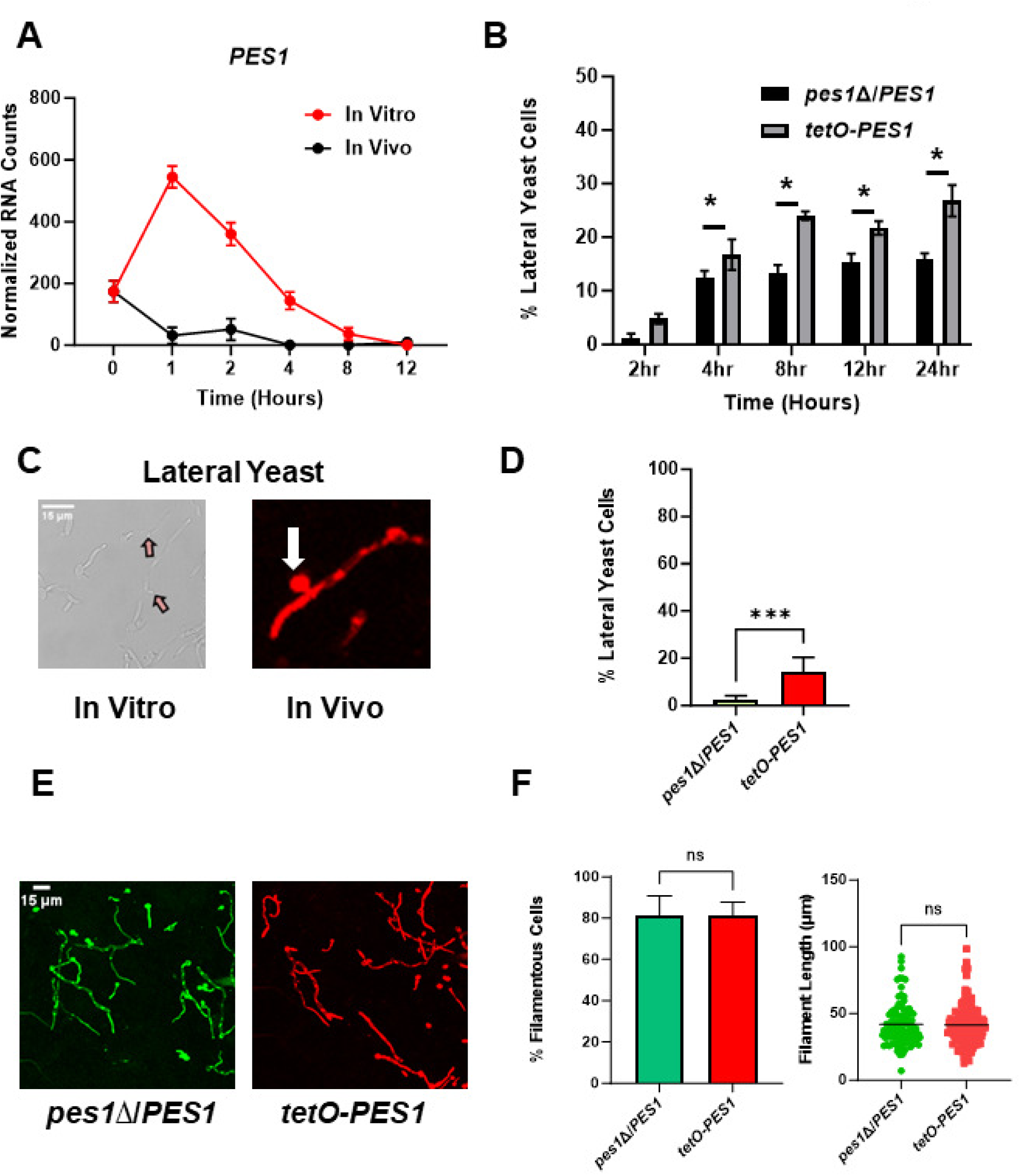
Heterologous expression of *PES1* is sufficient to induce lateral yeast formation in vivo. **A.** Normalized mRNA counts for the expression of *PES1*. Bars indicate normalized mRNA counts from three independent experiments with error bars indicating standard deviation. **B**. Effect of heterologous expression of *PES1* on lateral yeast formation over time course of in vitro hyphal induction. Bars indicate percentage of filaments with lateral yeast cells. * indicates statistically significant difference between groups (two-way ANOVA corrected for multiple comparisons with Sidak’s test). **C**. Representative images of lateral yeast in vitro and in vivo. **D**. Lateral yeast formation in vivo for *pes1*Δ/*PES1* and *tetO-PES1* 24hr post-infection. Bars indicate means from two independent experiments with standard deviation indicated by error bars. **** indicates p <0.0001 by Student’s t test. **E.** Representative images of the *pes1*Δ/*PES1* and *tetO- PES1* mutant morphologies in vivo. **F**. Quantification of % filamentous cells and filament length for the *pes1*Δ/*PES1* and *tetO-PES1* mutant strains. NS indicates no significant difference by Student’s t test (% filamentous cells) and Mann-Whitney test (filament length).

To determine if increased expression of *PES1* was sufficient to drive lateral yeast formation in vivo, we labelled the *tetO-PES1* and p*WT-PES1* strains with NEON and iRFP, respectively and inoculated a 1:1 mixture into the ear. In Fig. 7C, we show representative examples of lateral yeast identified in vivo which can be compared to hyphal branching with is shown in Fig. S1A. As shown in Fig. 7D, *tetO-PES1* strain formed dramatically more lateral yeast than the strain expressing *PES1* from the native promoter. This suggests that the low expression of *PES1* in vivo contributes to the near absence of lateral yeast and that increased *PES1* expression is able to drive lateral yeast formation in vivo. Interestingly, increased expression of *PES1* in vitro under some conditions reduces hyphae formation and the length of hyphae (15). However, the *tetO-PES1* strain formed hyphae to the same extent with similar lengths as the comparator strain (Fig. 7E&F).

Finally, reduced cAMP-PKA pathway activity has also been shown to increase the hyphae-to-yeast transition in vitro (26, 27). We found that mutants lacking the PKA pathway components Cyr1 and Tpk1/2 are able to form filaments in vivo (9). We, therefore, examined these strains for in vivo lateral yeast formation but found no significant difference relative control strains (Fig. S1B&C). For these experiments, we compared the *tetO-PES1/pes1Δ* strained to the corresponding *pes1*Δ/*PES1* heterozygous mutant. Therefore, it is possible that the heterozygous *pes1*Δ/*PES1* has lower lateral yeast formation due to haploinsufficiency (15). However, the lateral yeast formed by the *pes1*Δ/*PES1* mutant is not different than those formed by a SC5314-derived reference strain with both alleles of *PES1* (Fig. S1&C). Taken together, our data indicate that in the first 24hr of filamentation in vivo lateral yeast formation is extremely rare and that low *PES1* expression throughout the course of filamentation is likely to be a contributing factor.

## Discussion

Hyphal morphogenesis remains one of the most intensely studied aspects of *C. albicans* pathobiology (3). Most of this work has taken advantage of the wide range of in vitro media and conditions that induce the transition of *C. albicans* from a budding yeast morphology to the filamentous hyphal and pseudohyphae. We have begun to characterize this transition in mammalian tissue through the use of a in vivo imaging approach (6, 7. 8, 9). Here, we compared the time-dependence of in vivo filamentation and in the setting of a commonly used “host-like” induction condition (RPMI+10%BCS at 37°C). Although we have found some significant differences between in vivo and in vitro conditions, it is important to first point out the similarities, particularly during early time points. The rapid induction of positive regulators and repression of the major repressor of filamentation is observed under both conditions. By 2hr, there is significant overlap in the transcriptional responses based on a set of genes selected to include hyphae-related genes and environmentally responsive genes.

Generally speaking, recent studies have emphasized the fact that many features of hyphal morphogenesis are dependent upon the specific conditions used to induce that transformation (13). In the two conditions we studied, it appears that glucose availability is low in vitro. This conclusion is a based on the rapid increase in the expression of genes related to gluconeogenesis and alternative carbon metabolism in vitro but not in vivo. One possibility for this difference is that in vitro cultures have a fixed amount of glucose and other nutrients; once the fixed amount of glucose is depleted, the cells must switch to alternative carbon source metabolism. In vivo, *C. albicans* is in another organism that is delivering glucose and nutrients to its tissues and, as a result, delivering nutrients to *C. albicans*. After 12hr, genes such as that for alternative oxidase, *AOX2*, are beginning to be expressed at high levels, suggesting that the organism is experiencing reduced glucose delivery. This may be due to the beginning of abscess formation or tissue damage and necrosis, leading to disrupted blood flow in the region.

The most striking difference between these two conditions is that the expression of positive transcriptional regulators of hyphae morphogenesis and their targets declines sharply in vitro after a peak at the 4hr time point. This reduction in hyphae-associated gene expression correlates with a reduction in the filament-to-yeast ratio between 4hr and 12hr in vitro. The reduction in the filament-to-yeast ratio in vitro also correlates with the appearance of lateral yeast and follows a peak in the expression of the positive regulator of lateral yeast cell development, *PES1*. These observations suggest that the population of cells begins to express features of the hyphae-to-yeast transition after ∼4hr induction. In biofilm conditions, the lateral yeast cells lead to dispersion of yeast cells into the media and we suggest that the increase in lateral yeast cells between 2-4hr and the increase in planktonic yeast cells after 4hr may both be due to the hyphae-to-yeast transition (15, 25). Finally, the increase in lateral yeast formation in a strain that overexpresses *PES1* in vitro further supports the conclusion that this process is governed, at least in part, by this protein.

In vivo, we observed very little evidence of the hyphae-to-yeast transition during the time course examined. Furthermore, we have observed very few lateral yeasts in vivo up to 24hr post-infection and the filament-to-yeast ratios are stable over the same time period (7). Once again, these morphological observations can be correlated with the expression patterns of positive regulators of hyphae and the best characterized positive regulator of lateral yeast formation, *PES1*. Specifically, the expression of hyphae-induced transcriptional regulators (*BRG1*, *TEC1*, and *UME6*) increases over the first 1hr and then largely remains stable over the next 12hr and appears to maintain this pattern up to 24hr post infection based on previously reported data (7). This suggests that, in contrast to in vitro hyphae formation, the hyphal transcriptional program remains active throughout the first 12-24hr of in vivo infection.

*PES1* expression rapidly drops to near undetectable levels in vivo and remains low over the time course. Because lack of *PES1* expression inhibits lateral yeast formation in vitro (15, 25), this low expression seems likely to contribute to the low numbers of lateral yeast in vivo. As our data indicates, *PES1* expression from a heterologous promoter drives lateral yeast formation in vivo at a time point when it does not normally occur. Accordingly, this indicates that *PES1* expression is sufficient to trigger lateral yeast formation in vivo and strongly supports the conclusion that low expression of *PES1* is likely to contribute, at least in part, to the low rate of lateral yeast formation in vivo. This low level of *PES1* expression is not unique to the ear infection site and we have found similarly low levels of *PES1* expression in both infected kidney and tongue 24hr post-infection (7, 9, 28). Because of its role in virulence, it seems likely that its effects are most important after the initial establishment of infection. Indeed, Uppuluri et al. concluded that the role of *PES1* is most important after the establishment of infection based on the time dependence of its effect on fungal burden (29). Our observation of low expression of *PES1* in vivo is consistent with their findings and conclusions.

Our apparent inability to find strong evidence for the hyphae-to-yeast transition in vivo raises the interesting question: why not? A very simple explanation for the low rate of hypha-to-yeast transition in vivo may be that the transition may occur later during infection. The high expression of positive transcriptional regulators indicates that hyphal transcriptional program is consistently maintained though out the 24hr period we studied. As such, it seems very possible that the hyphal program begins to wane later in infection, as it does in vitro. Our ability to collect high resolution morphological data at time points beyond 24hr declines because *C. albicans* begins to form dense micro-abscesses in the ear tissue after 24hr (30).

In addition to providing insights into the dynamics of in vivo filamentation, our data have implications for the study of *C. albicans* filamentation in vitro. Specifically, the both the transcriptional and morphological features of the filamentation program in a given medium are likely to vary considerably over time, particularly in later time points. Therefore, it is important to be sure that the selected time point represents the specific stage of filamentation in which one is interested.

In summary, this works provides insights into the temporal dynamics of filamentation in mammalian tissue, allowing the identification of similarities and differences with a widely employed host-like in vitro conditions. Our previous work has found that there are significant differences in the sets of TFs (7) and protein kinases (9) that regulate in vivo filamentation relative to in vitro conditions. Many factors are likely to contribute to the differences in the regulatory factors required for in vivo filamentation. Based on this work, it seems likely that some of these differences could be due, at least in part, to the distinct environmental conditions encountered during infection and differences in the tempo of morphogenesis.

## Materials and methods

### Strains and media

The SC5314-derived *C. albicans* reference strain SN250 was used for all experiments except for those involving *PES1* mutant strains. The *pes1*Δ/*PES1* and *tetO-PES1* strains (15) were generous gifts of Julia Köhler (Harvard) and are in the SC5314 background. All *C. albicans* strains were precultured overnight in yeast peptone dextrose (YPD) medium at 30^0^C with shaking. Standard recipes were used to prepare YPD (31). RPMI 1640 medium was supplemented with bovine calf serum (10% vol/vol).

### In vitro hyphal induction

For in vitro hyphal induction, *C. albicans* strain was incubated overnight at 30^0^C in YPD media, harvested, and diluted into RPMI + 10% bovine calf serum at a 1:50 ratio and incubated at 37^0^C. Cells were collected at the different time points (e.g., 1hr, 2hr, 4hr, 8hr, and 12hr) and processed for microscopy or RNA isolation as described below.

### In vitro characterization of *C. albicans* morphology

Induced cells were fixed with 1% (v/v) formaldehyde. Fixed cells were then imaged using the Echo Rebel upright microscope with a 60x objective. The assays were conducted in triplicates on different days to confirm reproducibility.

### In vivo characterization of *C. albicans* morphology

These assays were performed as previously described (6). Briefly, 1 X 10^6^ WT *C. albicans* cells were inoculated intradermally in mouse ear (3 mice/time point). Mice (3 mice/time point) were sacrificed at each time points, and ears were harvested. One ear/mouse was immediately submerged into the ice-cold RNA later solution and another ear used for the imaging. A multiple Z stacks (minimum 20) were acquired and used it to score the yeast vs filamentous ratio. Round and/or budded cells were considered “Yeast”, whereas if the cells contain intact mother and filamentous which was at least twice the length of the mother body, were considered “filamentous.” Lateral yeast cells were distinguished from branching hyphae by requiring lateral yeast cells to be no more than two cell body lengths long and have curved than parallel cell walls (See Fig. 7C and Fig. S1A). A minimum of 100 cells from multiple fields were scored. Student’s *t* test was performed to define the statistical significance between the different time points. All animal experiments were approved by the University of Iowa IACUC.

### RNA extraction

In vitro and in vivo RNA extraction was carried out as described previously (7, 9). For in vitro RNA extraction, cells were collected at the different time points, centrifuged for 2 min at 10K rpm at room temperature and RNA was extracted according to the manufacturer protocol (MasterPure Yeast RNA Purification Kit). For in vivo RNA extraction, mice were euthanized, ear tissue harvested and the tissue placed into the ice-cold RNA-Later solution. Ear tissue was then transferred to the mortar and flash frozen with liquid nitrogen. Using pestle, the frozen was ground to the fine powder. The resulting powder was collected and 1 ml of ice-cold Trizol was added. The samples were placed on a rocker at RT for 15 min and then centrifuged at 10K at 4^0^ C for 10 min. The cleared Trizol was collected into 1.5 ml Eppendorf tube and 200 µl of RNase free chloroform was added to each sample. The tubes were shaken vigorously for 15 s and kept at RT for 5 min followed by centrifuge at 12K rpm at 4^0^ C for 15 min. The cleared aqueous layer was then collected to a new 1.5 ml Eppendorf tube and RNA was further extracted following the Qiagen RNeasy kit protocol.

### NanoString® gene expression analysis

NanoString analysis was carried out as described previously (7, 9). Briefly, in total, 40 ng of *in vitro* or 1.4 µg of *in vivo* RNA was added to a NanoString codeset mix and incubated at 65^0^ C for 18 hours. After hybridization reaction, samples were proceeded to nCounter prep station and samples were scanned on an nCounter digital analyzer. nCounter .RCC files for each sample were imported into nSolver software to evaluate the quality control metrics. Background subtraction was performed using negative control probes to establish a background threshold, which was then subtracted from the raw counts. The resulting background subtracted total raw RNA counts underwent a two-step normalization process. First normalized against the highest total counts from the biological triplicates and then to the wild type samples. Differentially expressed genes were defined as those with ± 2-fold change in expression with an FDR <0.1 as determined by Benjamini-Yeuketil procedure when compared to expression in the yeast phase inoculum which was grown overnight at 30°C in rich medium (yeast peptone with 2% dextrose, YPD).

### Software

Quantitative image analysis was carried out using ImageJ software. GraphPad Prism (V. 9.3.1) was used to plot the graphs and to perform the statistical tests.

## Acknowledgements

The authors thank Julia Köhler (Harvard) for providing *PES1* mutant strains. The authors also thank Scott Filler (UCLA) and Aaron Mitchell (Georgia) for helpful discussions and critical review of the manuscript. This work was supported by a National Institutes of Health Grant, R01AI133409 (DJK).

## Data Availability Statement

All raw (source) data and normalized data large-scale expression data generated by Nanostring are provided in Supplementary Tables 1 and 2. The bright field and confocal microscopy Z-stacks and images used to characterize *C. albicans* morphology at the different time points are very large files that are not easily deposited or annotated for deposit in public repositories. Therefore, the source data for the imaging figures are available from the corresponding author to interested investigators on reasonable request.

## Supplementary Tables and Figure

**Table S1.** Nanostring data for in vivo time course with raw counts, background corrected counts, normalized counts, fold change for each gene relative to either yeast phase, Student t-test values, and FDR calculated by the Benjamini-Yekutieli method. Differentially expressed genes were defined as those genes with ± 2-fold change in expression with FDR <0.1. Significantly upregulated genes are indicated by green fold change values at the time point in which they are differentially expressed; red indicates downregulated at that time point.

**Table S2.** Nanostring data for in vitro time course with raw counts, background corrected counts, normalized counts, fold change for each gene relative to yeast phase, Student’s test values, and FDR calculated by the Benjamini-Yekutieli method. Differentially expressed genes were defined as those genes with ± 2-fold change in expression with FDR <0.1. Significantly upregulated genes are indicated by green fold change values at the time point in which they are differentially expressed; red indicates downregulated at that time point.

**Supplementary Figure S1. A**. Representative images of hyphal branching in vitro and in vivo. Lateral yeast formation in vivo for *cyr1*ΔΔ (**B**) and *tpk1*ΔΔ *tpk2*ΔΔ (**C**) mutants 24hr post-infection. Bars indicate means from two independent experiments with standard deviation indicated by error bars. There were no significant differences (p> 0.05) between groups by Student’s t test.

